# Sexually antagonistic evolution of mitochondrial and nuclear linkage

**DOI:** 10.1101/2021.02.12.430938

**Authors:** Arunas Radzvilavicius, Sean Layh, Matthew D. Hall, Damian K. Dowling, Iain G. Johnston

**Affiliations:** Department of Mathematics, University of Bergen, Norway; Charles Perkins Centre, University of Sydney, NSW, Australia; School of Biological Sciences, Monash University, Victoria, Australia

## Abstract

Across eukaryotes, genes encoding bioenergetic machinery are located in both mitochondrial and nuclear DNA, and incompatibilities between the two genomes can be devastating. Mitochondria are often inherited maternally, and theory predicts sex-specific fitness effects of mitochondrial mutational diversity. Yet how evolution acts on linkage patterns between mitochondrial and nuclear genomes is poorly understood. Using novel mito-nuclear population genetic models, we show that the interplay between nuclear and mitochondrial genes maintains mitochondrial haplotype diversity within populations, and it selects both for sex-independent segregation of mitochondrion-interacting genes and for paternal leakage. These effects of genetic linkage evolution can eliminate male-harming fitness effects of mtDNA mutational diversity. With maternal mitochondrial inheritance, females maintain a tight mitochondrial-nuclear match, but males accumulate mismatch mutations because of the weak statistical associations between the two genomic components. Sex-independent segregation of mitochondria-interacting loci improves the mito-nuclear match. In a sexually antagonistic evolutionary process, male nuclear alleles evolve to increase the rate of recombination, while females evolve to suppress it. Paternal leakage of mitochondria can evolve as an alternative mechanism to improve the mito-nuclear linkage. Our modelling framework provides an evolutionary explanation for the observed paucity of mitochondrion-interacting genes on mammalian sex chromosomes and for paternal leakage in protists, plants, fungi, and some animals.

## Introduction

Sexual eukaryotes evolved in an ancient endosymbiosis between an archaeal host and the bacterial ancestors of modern mitochondria (Martin, 1998; Lane & Martin, 2010). Numerous mitochondrial genes have migrated to the nuclear genome and others have remained within the mitochondrial DNA (mtDNA) (Allen, 2003; Johnston & Williams, 2016). Key protein complexes of the mitochondrial respiratory chain contain subunits encoded by both mitochondrial and nuclear genomes (McKenzie et al., 2007), and mismatches between these two can be devastating to mitochondrial function and to cellular fitness (Rand et al., 2004; Wolff et al., 2014).

Mito-nuclear interactions are central to eukaryotic metabolism, and lifecycle traits that influence the linkage between interacting mitochondrial and nuclear alleles should experience strong selection. Many eukaryotes have evolved mechanisms that prevent the inheritance of paternal mitochondria (Greiner et al., 2015), and current theory views this maternal inheritance as a quality-control mechanism to improve selection against mitochondrial mutations (Hadjivasiliou et al., 2013; Radzvilavicius et al., 2017). However, low levels of paternal leakage of mitochondria often occur in protists, fungi, and many animals and plants (Greiner et al., 2015; Xu and Li, 2015), and this leaves the possibility that in some contexts nuclear alleles for paternal mtDNA transmission may be advantageous.

The strength of statistical associations between nuclear and mitochondrial genes differs across the two sexes, and these differences may give rise to sex-specific fitness effects and sexual conflict (Radzvilavicius et al., 2017). Theoretically it may be expected that mtDNA mutational diversity will have male-detrimental fitness effects, because-male specific nuclear loci evolve largely independently of the mitochondrial background (Frank and Hurst, 1996; Gemmell et al., 2004). Empirical evidence of strong sex-specific effects of mtDNA diversity in natural populations is scarce (Beekman et al., 2014; Dowling and Adrian, 2019), and how organisms escape these sex-specific fitness effects is poorly understood (Wade and Brandvain, 2008; Beekman et al., 2014; Kuijper et al., 2015).

In the nucleus, recombination and random chromosome assortment will also change the extent of statistical associations between the mitochondrial and nuclear genomic components. Tight linkage to female sex determination will increase the associations when there is maternal mtDNA transmission, and recombination between the two will reduce them. In males, we expect to see the opposite linkage pattern of weak mito-nuclear linkage for alleles segregating with male sex determination. Theory predicts that female sex chromosomes should be enriched in mitochondria-interacting genes to increase co-transmission of mitochondrial and nuclear gene combinations (Brandvain et al., 2007; Brandvain and Wade, 2009). Observations, however, suggest that there either is no bias in gene distribution, or that mitochondrial genes are underrepresented on the X chromosome (Drown et al., 2012; Dean et al., 2014; Rogell et al., 2014). Our theoretical understanding of how selection acts on linkage patterns among the mitochondrion-interacting genes in the nucleus is therefore incomplete.

These divergent patterns of mito-nuclear linkage arise because of gamete fusion (syngamy) with female-biased mitochondrial transmission, and, more generally, they are the consequence of sex. Most eukaryotes can reproduce sexually, and this widespread occurrence suggests that sex has a universal selective advantage over obligately clonal reproduction. Theory of meiotic sex evolution focuses on fitness effects of breaking down nuclear linkage disequilibria that build up through selection, mutation, and drift (Kondrashov, 1993). These models revealed that the evolution of recombination often requires special conditions and specific gene interaction nonlinearities that are not common in eukaryotes (Otto, 2007). Eukaryotes are both ancestrally sexual and endowed with mitochondria (Ramesh et al., 2005; Malik et al., 2008), and it is therefore possible that the ancient evolution of sex was the result of the fitness effects associated with mtDNA mutational diversity and the complex interplay between mitochondrial and nuclear genomes (Lane, 2011; Havird et al., 2015; Radzvilavicius, 2016).

In this work our aim is to understand the evolution of nuclear regulators of genomic linkage between sex determination, mitochondrial and nuclear alleles, and to understand how the evolution of these linkage patterns differs across sexes, that is, sexual antagonism of linkage evolution. We propose that linkage disequilibria arising with maternal mtDNA inheritance may select for sex-independent segregation of nuclear loci interacting with mitochondria and for paternal leakage of mitochondria, and that recombination may reduce the expected male-harming fitness effects. To test this hypothesis, we develop a novel modelling framework describing nuclear-mitochondrial population genetics with nuclear alleles regulating the linkage associations between mitochondrial and nuclear alleles: recombination within the nucleus and paternal leakage of mitochondria. Fitness is determined by how well mitochondrial haplotype distribution within the cell matches the nuclear background. The model reveals strong sex-specific fitness effects of sex-linkage of mitochondria-interacting loci and paternal leakage: males evolve high recombination rates improving their mito-nuclear match and females evolve to suppress independent segregation in the nucleus. Recombination evolves in both sexes when meiosis regulation is not strictly linked to sex determination, and paternal leakage of mitochondria can evolve in both sexes when it improves the mito-nuclear linkage.

## Model summary

We model nuclear-mitochondrial population genetics, with a haploid nucleus and *N* mitochondria per cell (see Methods). Nuclear alleles, generally loosely linked to the mitochondrial populations, regulate mitochondrial diversity, the mode of mtDNA inheritance (ranging from strict maternal inheritance to biparental mitochondrial mixing), and recombination between sex-determination and mitochondria-interacting nuclear loci. Sex (male or female) is determined at the haploid stage, and mitochondria are inherited predominantly from the female gamete. Two mitochondrial haplotypes segregate within the population, *a* and *A*, with the mutation rate between the two *μ*, and we model a single nuclear locus which interacts with mitochondria. The nuclear locus has two allelic variants *b* and *B*, well matched either to the haplotype *a* or *A*, with the nuclear mutation rate between the two versions *v*. Cell fitness depends on the total number of mismatches between the two genomes *n* : *w*(*n*) = 1 − *s*(*n*⁄*N*)^*x*^, where s is selection strength, and we set *x* = 2 to model the so-called mitochondrial threshold effects, whereby functionality depends on mutant proportion in a non-linear way – the deleterious effect of each new mutation becomes stronger as fewer well-matched mitochondria are left to compensate (Rossignol et al., 2003; Johnston & Burgstaller, 2019). More explicitly, the fitness of a cell with *i* mitochondria of haplotype *a* and the nuclear allele *B* is 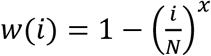, while the fitness of a cell with the alternative nuclear allele *b* (and *i* mitochondria of type *a*) is 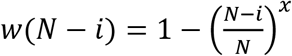.

Each generation, organisms experience the life cycle of 1) mitochondrial and nuclear mutation, 2) haploid selection, and 3) reproduction, which can be either clonal or sexual. In sexual reproduction, haploid cells of opposite mating types fuse at rate *r*_*sex*_. Sexual cell fusion in our model is present independently of recombination, and it may have evolved as a first step towards fully sexual life cycles (Radzvilavicius, 2016b). Because of the uniparental inheritance of mitochondria, all *N* mitochondria of the maternal mating type are transmitted to the zygote, while the paternal gamete transmits only *lN*, where paternal leakage *l* ≤ 1. The resulting diploid zygote can undergo recombination between sex-determination locus and *b*/*B* locus with probability *r*, which is followed by reductive cell division in which mitochondria are randomly distributed into two daughter cells. Cells can alternatively reproduce clonally (rate 1 − *r*_*sex*_), in which case the mitochondrial contents of the cell are doubled and then distributed randomly across the two daughter cells.

Meiotic recombination is regulated by nuclear loci that segregate together with sex-determination locus (we later consider some recombination between the two). This model structure allows us to study the fitness effects of recombination and its evolution in the two sexes independently. We expect these sex differences to be important because of the female-biased mitochondrial inheritance, which will create sex-specific linkage disequilibria. First, we fix model parameters and study the sex-specific fitness effects of recombination, paternal leakage and sex rates. Then, we model the evolution of recombination between sex-determination and mitochondria-interacting nuclear loci, and the evolution of nuclear alleles regulating paternal leakage.

## Results

We first sought to understand the relationship between the mito-nuclear match and fitness, and the life cycle parameters (recombination rate *r*, paternal leakage *l* and sex rate *r*_*sex*_) in populations where nuclear and mitochondrial mutations are at equilibrium with purifying selection. With strict maternal inheritance of mitochondria (paternal leakage *l* = 0), mtDNA is transmitted exclusively through the maternal line. Maternal mitochondria are always inherited together with the maternal nuclear loci, while male-specific nuclear loci receive a new mitochondrial environment every sexual generation. Because of these differences in the strength of inter-genomic associations when there is no recombination, selection maintains a strong mito-nuclear match in females, while males accumulate mismatch mutations (Fig. 1a).

**Figure 1.**
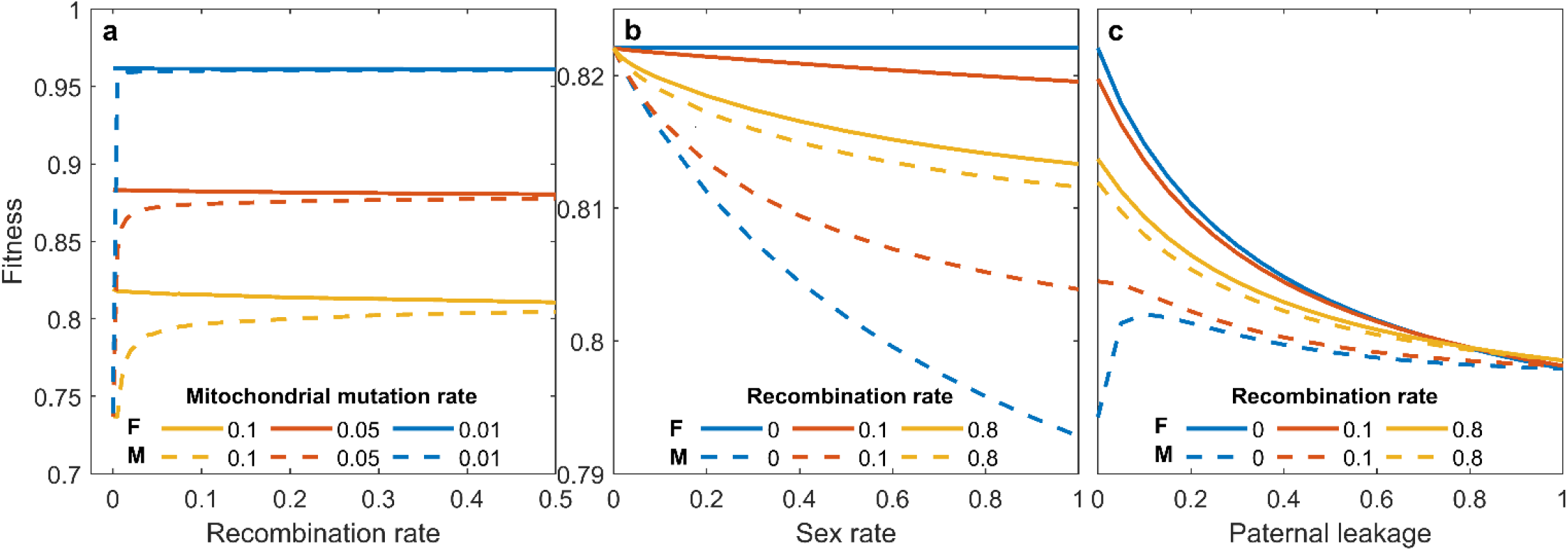
Sexual dimorphism in fitness effects of mito-nuclear mismatch. (a) Males (M), accumulate mitochondrial mismatch mutations more rapidly than females (F), impairing fitness. Recombination between male and female nuclear loci improves the match, restoring male fitness. *v* = 0.01, sex rate *r_sex_* = 0.9, *l* = 0. (b) Fitness differences between females and males are most striking with high cell fusion rates *r_sex_*. Recombination increases male fitness with frequent sexual cell fusion but has little effect with rare sex. *μ* = 0.1, *v* = 0.01, *l* = 0. two sexes for *r* = 0.5 (yellow lines). (c). If there is no meiotic recombination, small amounts of paternal leakage improve the mito-nuclear match in males, also improving fitness. Paternal leakage always reduces female fitness, as it erodes mutational variance in mitochondrial populations that has built up through segregation. *μ* = 0.1, *v* = 0.01, *r_sex_* = 0.9.

When there is recombination, mitochondrion-interacting loci in both male and female offspring may be transmitted together with maternal mitochondria. In males, this type of transmission improves linkage and increases the efficacy of selection for mito-nuclear match (Fig. 1a). Recombination reduces the linkage strength in females, but this has a much weaker influence on their fitness, as the relative fitness effect of mito-nuclear mismatches is initially weak and increases with each new mismatch mutation because of the mitochondrial threshold effects (Rossignol et al., 2003).

Differences in mito-nuclear linkage associated with recombination also depend on the rate of sexual cell fusion *r*_*sex*_. With obligate sex, selection among males acts on new mitochondrial-nuclear gene combinations every generation, and when sex is rare, linkage persists for multiple generations. This change is reflected in male fitness relative to females, which is highest with low rates of sex and declines as sex becomes more frequent (Fig. 1b).

Finally, because the inheritance of mitochondria is not always strictly maternal, paternal leakage *l* will alter the strength of mito-nuclear linkage in both sexes. With *l* > 0, a fraction of paternal mitochondria is transmitted together with male nuclear loci, increasing the linkage, and in females the effect is the opposite. Paternal leakage also increases the extent of heteroplasmy and reduces variance among cells, reducing the intensity of selection against mito-nuclear mismatches (Radzvilavicius et al., 2017). In balance, paternal leakage is always detrimental to females (Fig. 1c). In males, small amounts of paternal leakage increase equilibrium fitness, but only if there is zero or little recombination among the nuclear loci.

## Evolution of recombination

We have seen that both recombination and paternal leakage of mitochondria influence the strength of statistical associations between mitochondrial and nuclear genomes, inducing diverging fitness effects in males and females. Next, we asked if these fitness effects can select for the spread of nuclear mutations reducing linkage in the nucleus through recombination between sex determination and mitochondrion-interacting loci.

Our model reveals that mito-nuclear interactions can maintain the joint polymorphism in mitochondrial and nuclear genotypes (Fig. 2a-d), but the mitochondrial haplotype distributions differ between the two sexes. When there is no paternal leakage, female mitochondrial haplotypes match their nuclear backgrounds. Males inherit mitochondria from females with both nuclear alleles irrespective of their own nuclear background, and this generates frequent mito-nuclear mismatches (Fig. 2a), explaining their reduced fitness in Fig. 1. When overall haplotype distributions are biased towards one allelic variant (Fig. 2c), the more common nuclear allele also becomes overrepresented among males (Fig. 2d). With recombination, male mitochondrial distributions approximate the haplotype distributions in females, restoring the match and improving fitness (Fig. 2a, c).

**Figure 2.**
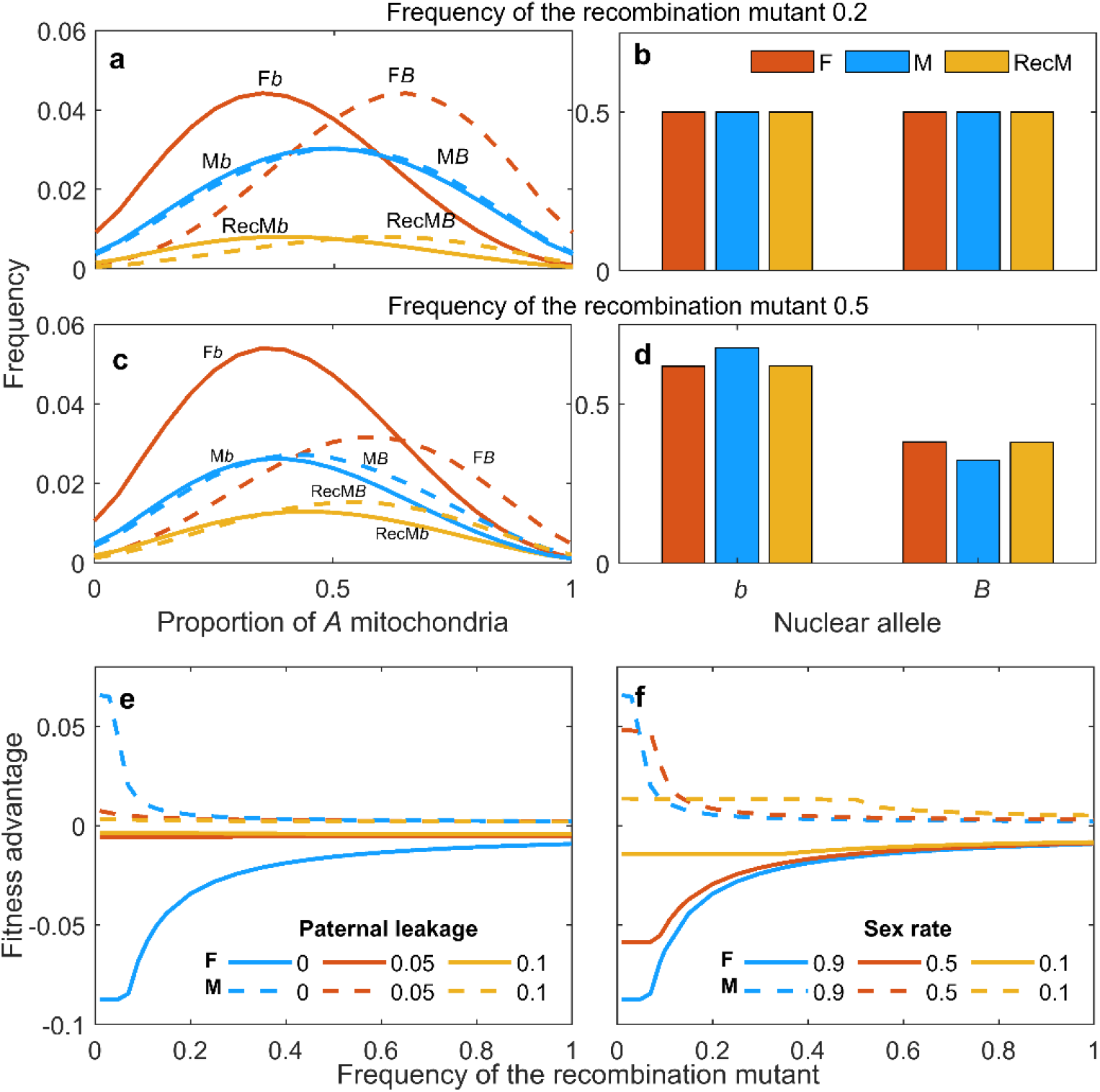
Evolution of recombination in the two sexes. (a-d) Mitochondrial (a, c) and nuclear (b, d) allele distributions across the two sexes. In females (red, F), mitochondrial haplotypes match the nuclear background (*b* or *B*), and in males (blue, M) there is little difference in mitochondrial distribution across the two nuclear alleles. Recombination in males (yellow, RecM) improves the mito-nuclear match, but there is little change in the distribution of nuclear alleles *b* and *B* (b). Frequency of recombining males in (a) and (b) is 20%. Mitochondrial mutation rates in this figure are symmetrical for illustrative purposes, but strict symmetry is not a requirement for the polymorphism. High frequencies of recombinant males (50% in (c) and (d)) break the symmetry in haplotype distributions and the dominant nuclear allele *b* becomes overrepresented in males (d, blue) because of the maternal mtDNA inheritance. (e, f) Fitness advantage of the recombination mutant (*r* = 0.5) in an ancestral population of non-recombining males (M) and females (F). The recombination regulation locus segregates together with sex-determination. Because of its effects on mito-nuclear linkage across the two sexes, meiotic recombination is expected to evolve in males but not in females. Paternal leakage (e) and rare facultative sex (f) both reduce the fitness advantage of rare recombination in males. *μ* = 0.1, *v* = 0.01, *r_sex_* = 0.9 in (e) and *l* = 0 in (f). *μ* = 0.1, *v* = 0.05, *r_sex_* = 0.9, *l* = 0 (a-d)

We next studied recombination-inducing mutations (inducing recombination at a rate *r* = 0.5) arising in a population of sexually fusing cells that do not recombine (*r* = 0). These new mutations arise linked to the sex determination locus, and initially there is no recombination between the two. The spread of these mutations is determined by their fitness advantage over the non-recombining subpopulations (Fig. 2e, f). The fitness advantage associated with the recombining mutant is always positive in males, while female recombination mutants are at disadvantage. In males, low amounts of paternal leakage (Fig. 2e) and frequent sex (Fig. 2f) both promote the evolution of recombination when it is rare, and in females we find the opposite evolutionary pattern.

Alternatively, sex-determination and recombination-regulation loci may be segregating independently, and meiotic segregation of nuclear genes is then regulated by both sexes. We find that recombination-inducing mutations can evolve in both females and males, although their frequencies are generally higher in males (Fig. 3). Recombination mutants reach high frequencies when the genomic distance between sex-determination and meiosis-regulation is short, when paternal leakage is rare (Fig. 3a) and when sex is infrequent (Fig. 3b), as all these effects increase the linkage of the recombination-regulation locus to male mitochondrial and nuclear genotypes that initially experience its positive fitness effects.

**Figure 3.**
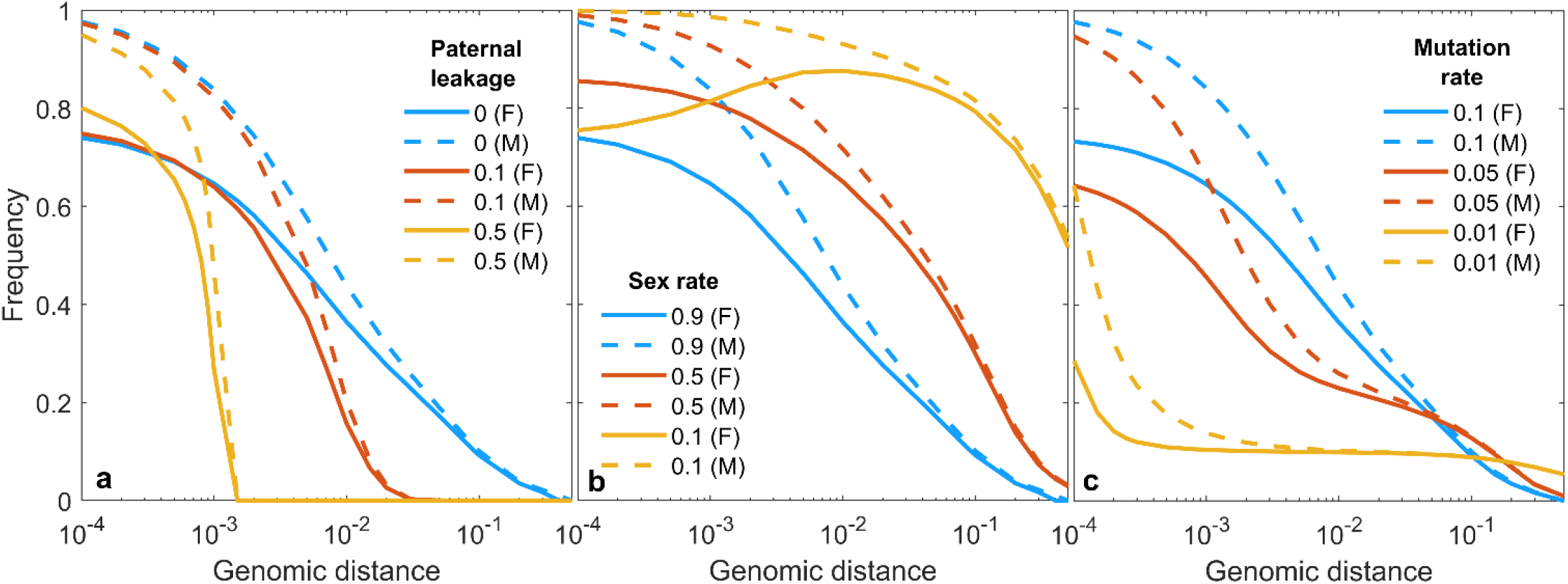
Evolution of meiotic recombination without tight linkage to sex determination. The equilibrium frequency of recombination-inducing mutant (*r* = 0.5) in the two sexes (F – females, M – males) depends on the genomic distance between the sex-determination and recombination-regulation loci. Crossover-inducing mutations evolve in both sexes with rare recombination between the two loci, but not when recombination regulation locus segregates independently of sex determination. When there is paternal leakage, recombination evolves to a lower frequency (a). Infrequent sex creates strong linkage associations between the regulator allele and mitochondrial populations, and this increases the frequency of recombination allele (b). High mtDNA evolution rates select for more recombination in the nucleus (c). *μ* = 0.1 in (a) and (b), *v* = 0.01, *r_sex_* = 0.9 in (a) and (c) and *l* = 0 in (b) and (c).

High mitochondrial mutation rates select for more common recombination in the nucleus (Fig. 3c). This is because fast mitochondrial evolution maintains the stable mito-nuclear polymorphism within the population. When mtDNA mutation rates are low, this joint polymorphism is more likely to collapse and then there is little fitness advantage to recombination. The strong mitochondrial haplotype polymorphism is also more likely to collapse when recombination itself becomes frequent, as evident from the rapid decline in the fitness advantage of recombination mutants in Fig. 2e-f.

## Evolution of paternal leakage

We have seen that when recombination is rare, paternal leakage can increase the strength of mito-nuclear linkage and improve fitness (Fig. 1). Kuijper et al. (2015) have suggested previously that low amounts of paternal leakage can eliminate the evolution of detrimental sex-specific fitness effects. We therefore asked if paternal leakage, regulated by a nuclear locus in either females or males, can evolve as an alternative to recombination of mitochondrion-interacting loci when fitness is determined by how well the mtDNA populations match the nuclear background.

Nuclear mutations increasing the extent of paternal leakage alter the strength of statistical associations between the nuclear genotype and the mitochondrial haplotypes, and we find that they can be advantageous in both males and females (Fig. 4a, b). Because of the predominantly maternal inheritance of mitochondria, this statistical linkage is stronger in females, and if recombination is infrequent, paternal leakage will evolve to 0 to maintain this strong linkage. When recombination rates are high, female nuclear alleles that are well adapted to their mitochondrial populations may be replaced by male copies, and this weakens the mito-nuclear linkage. Females therefore evolve paternal leakage only when sex and meiotic recombination are frequent (Fig. 4c). Male nuclear alleles evolve to increase paternal leakage when recombination is rare (Fig. 4d), because biparental mtDNA transmission increases the odds that nuclear and mitochondrial alleles will be inherited together, and selection can then act to improve the mito-nuclear match over generations.

**Figure 4.**
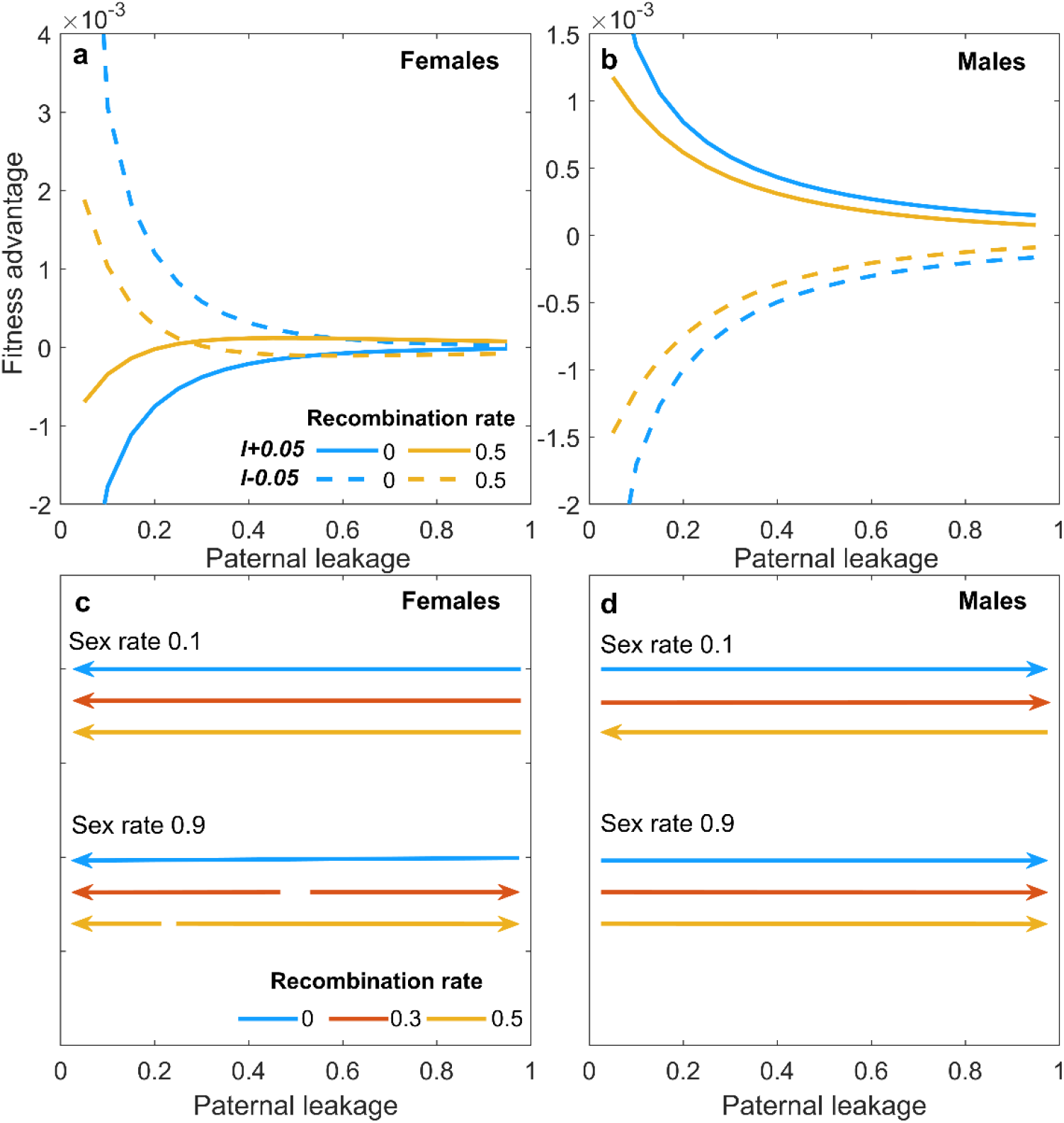
Evolution of paternal leakage regulated by nuclear alleles. (a, b) Fitness advantage of rare nuclear mutations increasing or reducing the amount of paternal leakage *l* by 1/*N* = 0.05 (*l* + 0.05 and *l* − 0.05) when paternal leakage is regulated by females (a) and males (b). *r_sex_* = 0.9, *v* = 0.01. (c, d) Direction of paternal leakage evolution in females (c) and males (d) depends on sex and recombination rates. Females evolve high rates of paternal leakage only when both sex and meiotic recombination are frequent, as only then biparental mtDNA inheritance can improve mito-nuclear linkage. Males evolve to increase paternal leakage when sex is frequent and when recombination rates are low, because then the linkage between their mitochondrial and nuclear genotypic components is weak.

## Discussion

Mitochondria play vital bioenergetic roles in eukaryotes, and mtDNA works in tandem with the nuclear DNA to regulate the most central aspects of mitochondrial energy production. Disrupted mito-nuclear match can have severe fitness consequences (Burton et al., 2006; Reinhardt et al., 2013; Ma et al., 2016; Dobler et al., 2018; Giordano et al., 2018). We therefore expect that eukaryotic life-history and reproductive traits that influence the linkage associations within and between the two genomes will experience strong selection. Can these linkage effects help explain the evolution of meiotic recombination in ancestral eukaryotic lineages?

Our model revealed sexual conflict in mito-nuclear linkage evolution. Because of the predominantly maternal mitochondrial inheritance, the genetic linkage between sex-specific nuclear loci and mitochondria is much stronger in females than it is in males, where an unrelated mitochondrial population is inherited every sexual generation. Males therefore tend to accumulate more deleterious mutations in mitochondria-interacting loci, and there is selection to increase the mito-nuclear linkage in males. We found that when loci regulating nuclear linkage segregate in tight linkage with sex determination, males evolve recombination of mitochondria-interacting loci, and females evolve to increase linkage to their sex determination locus. Drown et al. (2012) have previously proposed that sexual conflict may be responsible for their observations of the relative scarcity of mitochondria-interacting genes on the mammalian X chromosome, co-transmitted with mtDNA at a higher rate than autosomes. Empirical support of these predictions comes from observations in *Drosophila*, where nuclear-mitochondrial gene duplicates tend to relocate from the X to autosomes (Galach et al. 2010). We expect that mito-nuclear interactions will select for this type of sex-independent segregation of mitochondrion-interacting loci in males, and in both sexes if the linkage between the two is regulated by nuclear loci that themselves are not sex-specific.

Selection for the mito-nuclear match also creates sexual conflict in paternal leakage regulation. When sex is frequent, male nuclear alleles will evolve to increase the rate of paternal leakage, and female-specific nuclear alleles will evolve to reduce leakage when mitochondria-interacting loci are linked to sex-determination loci. In our previous work we found that even without direct mito-nuclear interactions the diverging patterns of linkage in males and females may create sexual conflict over mitochondrial inheritance regulation, and that males may select for paternal leakage (Radzvilavicius et al., 2017). Together, these results may explain some of the variation in mtDNA transmission modes observed in plants, fungi and many animals that have low or moderate levels of paternal leakage of mitochondria and plastids (Woloszynska, 2010; Greiner et al., 2015; Xu and Li, 2015). We saw that mito-nuclear interactions can maintain a stable mitochondrial haplotype polymorphism within the population, but that polymorphism tends to collapse when there is recombination or paternal leakage. These observations establish a link between the mechanisms regulating mitochondrial and nuclear genetic diversity at the level of the cell, and empirically observable population-level haplotype distributions.

Evolutionary theory predicts that mutational variability in maternally transmitted mtDNA may have deleterious effects in males, because male-specific nuclear alleles evolve independently of the maternally inherited mitochondrial background (Frank and Hurst, 1996; Gemmell et al., 2004). Sex-biased effects of mitochondrial diversity have been observed in experiments (Vaught and Dowling, 2018; Dowling and Adrian, 2019), but the evidence in natural populations is scarce (Beekman et al., 2014). This could be because mitochondrial mutations in fact respond to selection in males, as population-genetic models predict would happen when there is inbreeding or kin selection (Wade and Brandvain, 2009; Keaney et al. 2019), or when both parents contribute their mitochondria to the zygote (Kuijper et al., 2015). Our current results show that both paternal leakage and reduced sex-linkage of nuclear genes interacting with mitochondria may evolve to reduce the severity of these predicted male-harming effects.

We believe that selection acting on mito-nuclear linkage associations may have been one of the main forces responsible for the evolution of sexual life cycles with syngamy and meiotic recombination in ancestral eukaryotes. While virtually all theoretical models developed thus far have focused on genomic linkage between interacting loci within the nucleus (Kondrashov, 1993), the most unique aspects of eukaryotes related to the mitochondrial endosymbiosis and the ensuing genomic transformation, new types of genetic interactions between the two cellular compartments and mitochondrial diversity effects, are all missing from this theory. Eukaryotes are the product of an ancient endosymbiosis between two types of prokaryote – an archaeal host and bacterial ancestors of mitochondria (Martin, 1998). Prokaryotes reproduce asexually with frequent horizonal gene transfer, but almost all eukaryotes are sexual and all evolved from a sexual ancestor. Why did the ancient lineages of emerging eukaryotes evolve sexual cell fusion and meiotic recombination in their nuclear chromosomes, traits that are completely missing in prokaryotes? We contend that the answer lies in the ancient endosymbiosis that produced the eukaryotic cell.

Previously, we have shown that increasing intracellular mtDNA diversity may have selected for the evolution of syngamy in populations without sexes or mating types (Radzvilavicius, 2016). Because of mitochondrial threshold effects, mitochondrial mixing between vertical eukaryotic lineages has an average positive fitness effect, because it reduces the odds of inheriting an unusually high, above-threshold number of mtDNA mutations. If the ancient eukaryotes did not have mechanisms regulating the number of mitochondrial endosymbionts, syngamy would have also helped maintain a constant number of bioenergetic organelles, and there may have been other ecological conditions that favored the increased intracellular mtDNA diversity. The evolution of syngamy would have altered the linkage associations between mitochondrial and nuclear genes, the effect exacerbated in an asymmetric way by the evolution of uniparental mitochondrial inheritance. We have shown that the asymmetry in how selection acts on mitochondrial-nuclear associations may have selected for the evolution for recombination among nuclear genes to improve the mito-nuclear match in the mating type that contributes fewer mitochondria to the zygote. Future work should explore the alternative ways in which mitochondrial-nuclear interactions can select for the evolution of meiotic crossover in genomic lineages that are already independent of mating type or sex-determination systems, for example, to facilitate the evolution of nuclear alleles that compensate for the negative effects of mitochondrial Muller’s Ratchet, as suggested by Havird et al. (2015).

## Data availability

MATLAB/Octave scripts implementing the model are available at https://doi.org/10.5281/zenodo.4537142.

## Acknowledgements

This project has received funding from the European Research Council (ERC) under the European Union’s Horizon 2020 research and innovation programme (Grant agreement No. 805046 (EvoConBiO) to IGJ). Arunas Radzvilavicius thanks Nicky Gluch (Sydney), Josh Christie (Sydney), and Kostas Giannakis (Bergen) for their valuable and honest feedback.

## Methods

Here we describe the transition-matrix methodology for the joint nuclear-mitochondrial population genetics, first introduced in Radzvilavicius, 2016. This framework represents genotype frequency distributions in infinite populations as matrices ***P***, and the time evolution of the P-matrices is then modelled using multiplication by a set of square transition matrices ***Y***_***n***_, ***P***(next) = ***Y***_***n***_ ***P***(current), where each of the transition matrices ***Y***_***n***_ is constructed to represent probabilistic life-cycle events: mutation, selection and reproduction. The explicit form of these matrices is described below. This framework gives exact changes in genotype frequency distributions between life cycle events and between generations (Radzvilavicius et al., 2017). In practice, we initialize all P-matrices with random mutational distributions and track their exact evolution using numerical linear algebra methods.

We model infinite populations of unicellular eukaryotes, each with *N* mitochondria and a haploid nucleus. There are two mitochondrial haplotypes segregating within the population, *a* and *A*, and a nuclear locus *b*/*B* interacting with mitochondria. The second nuclear locus *h*/*H* regulates the evolving lifecycle parameter, and the third haploid locus determines sex. We represent genotype frequency distributions within males and females using *N* + 1-by-*K* matrices ***P***^***sn***^ (one matrix for each sex *s*, ***P***^***fn***^ and ***P***^***mn***^, because we do not model deviations from 1:1 sex ratio), where *K* is the number of possible nuclear genotypes and *N* + 1 is the number of possible mitotypes. Matrix element 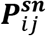denotes the frequency of individuals with *i a*-type mitochondria (row index) and (*N* − *i*) *A*-type mitochondria, and a nuclear allele at the *b*/*B* locus *j* (*j* = 1 (*B*), *j* = 2 (*b*)), and the allele at the *h*/*H* locus *n*. Each generation, the population goes through the life cycle of 1) mitochondrial and nuclear mutation, 2) haploid selection, and 3) reproduction, which can be either clonal or sexual.

### Mutation

Genotype distribution frequencies after mitochondrial and nuclear mutation become

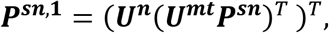

where the superscript 1 denotes the lifecycle stage after mutation. The mitochondrial mutation matrix ***U***^***mt***^ is a *N* + 1 square matrix and the matrix element 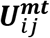 denotes the probability that a cell with *j a*-type mitochondria will have *i a*-type mitochondria after the bidirectional mutation from *a* to *A* (mutation rate *μ*_2_) and from *A* to *a*:

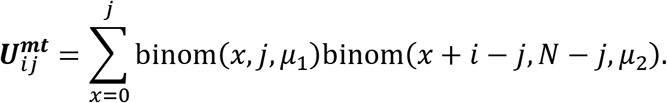

The binomial probability distribution function binom 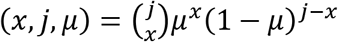 represents the probability of *x* successes in sampling with replacement from the population of size *j*, each with success probability *μ*. We sum over *x* counting all possible mutant number combinations from *a* to *A* and back from *A* to *a* that add up to *i* − *j*.

The nuclear mutation matrix element 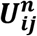 denotes the transition probability from *j*-th to *i*-th nuclear genotype (a or *A*) with the symmetric mutation rate *v*:

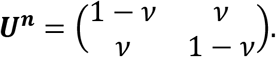

### Selection

Selection in the model acts at the haploid phase of the life cycle, after the mutation accumulation stage. Cell fitness depends on the total number of mismatches between the two genomes and can be expressed as *N* + 1-by-2 matrix ***W***, with elements 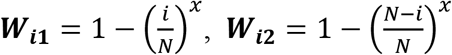, where *x* > 1 accounts for the mitochondrial threshold effects. The first column of ****W**** represents the nuclear allele *B*, and the number of mito-nuclear mismatches is represented by the row number *i* (the number of *a*-type mitochondria. The second column of ****W**** corresponds to the nuclear allele *b*, and the number of mismatches is then *N*-*i*.

The selection step resamples genotype frequencies weighted by their relative fitness, and mito-nuclear genotype frequencies after selection then become

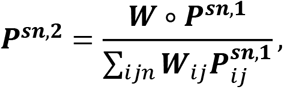

where (∘) denotes matrix Schur product, and the superscript 2 denotes the lifecycle stage after selection.

### Reproduction

In the next step, organisms undergo sexual cell fusion with probability *r*_*sex*_, or divide clonally otherwise. In clonal reproduction we duplicate the mitochondrial population of each cell and then segregate it randomly among the two daughter cells: ***P***^***sn*,3*asex***^ = ***AP***^***sn***,**2**^, where resampling matrix ***A*** has elements that follow the hypergeometric distribution,

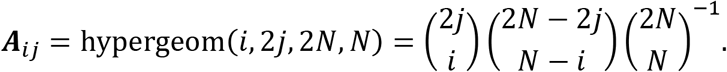

If the reproduction is sexual, pairs of cells of opposite mating types fuse, mixing their mitochondrial contents according to the value of paternal leakage *l*. The diploid zygote has *N* maternal mitochondria and *lN* paternal mitochondria sampled from the father without replacement. There are 16 diploid nuclear genotypes, and their joint mito-nuclear genotype frequency distributions are

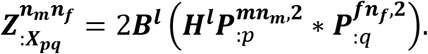

Here 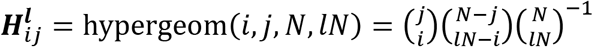, which we include to resample the number of mitochondria donated by the paternal gamete from *N* to *lN* without replacement. Transition matrix ***B*** has binomial elements 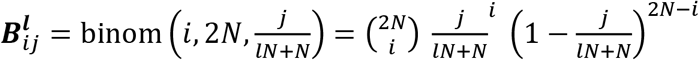, and is included to restore the number of zygote mitochondria back to the full diploid complement of 2*N* sampling with replacement. The index notation : *p* denotes the *p*-th column of the given matrix, and an asterisk denotes vector convolution, 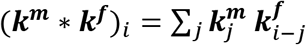, as the number of mitochondrial mutants in the zygote *i* represents all possible combinations of mutant numbers in the two gametes that add up to *i*. The nuclear genotypes indexed as *p* and *q* come from male and female gametes, respectively. Matrix 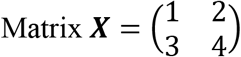 maps haploid nuclear indices *p* and *q* into diploid nuclear genotype index ***X***_*pq*_.

Cell fusion is followed by nuclear recombination between sex-determination locus and *b*/*B* locus with probability *r*. Recombination operator ***R*** consists of transition probabilities between diploid nuclear genotype index pairs ***X***_*pq*_. After recombination,

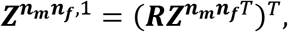

where recombination transition matrix is

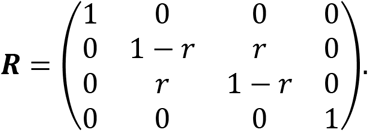

Recombination is followed by two reductive cell divisions back into the haploid phase,

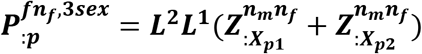

for females and

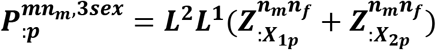

for males. At the start of the next generation the mito-nuclear genotype frequencies are

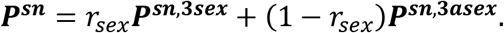

### Numerical methods

To study mean male and female fitness at mutation-selection equilibrium 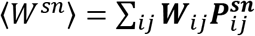, we solved the system of matrix equations for the steady-state P-matrices with only wild type nuclear alleles *n* regulating the lifecycle parameter. This equilibrium is defined by the self-consistency requirement, where at generation *k*, ***P**^**sn**^*(*k* − 1) = ***p^sn^***(*k*), and it is independent of initial conditions. We solve for self-consistency using fixed-point iteration with random initial conditions with the numerical accuracy of 10^−10^. This means that all genotype frequencies stay constant between generations within the range of 10^−10^.

Next, we model mutations in nuclear loci regulating the lifecycle parameter (recombination rate *r* or paternal leakage *l*) that arise at frequency *f* in generation *k* + 1:

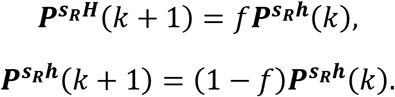

We then either kept the frequency *f* constant and measured the fitness advantage of the mutant 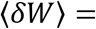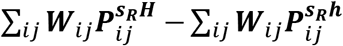 at equilibrium (Fig. 2e and 2f, Fig. 4a and 4b), or tracked the frequency of the mutant 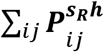 until it reached equilibrium (Fig. 3).

To produce Figure 4, for each possible value of wild-type paternal leakage *l* we studied the evolution of paternal leakage mutants *L* = *l* ± 1 ⁄ *N* (where *N* is the total number of mitochondria in the cell, and the smallest difference in paternal leakage corresponds to a single paternal mitochondrion). Because these are mutations with weak fitness effects, their fitness advantage is approximately independent of their frequency. Therefore, they either uniformly increase in frequency and replace the wild-type allele, or their frequency decreases towards zero. Although the fitness advantage of the new mutants is frequency-independent, we used the initial frequency of *f* = 0.01, and then tracked the fate of the mutant as it evolves to the frequency of 1 (replacing the wild-type allele) or the frequency of 0. The direction of an arrow in Fig. 4 represents mutant replacement and the opposite direction represents stability against new mutations in paternal leakage.

## Notes

https://doi.org/10.5281/zenodo.4537142

### Competing Interest Statement

The authors have declared no competing interest.

## References

Allen JF. 2003. Why Chloroplasts and Mitochondria Contain Genomes. Comp. Funct. Genomics 4(1), 31–36.

Beekman M, Dowling DK, Aanen DK. 2014. The costs of being male: are there sex-specific effects of uniparental mitochondrial inheritance? Philos. Trans. R. Soc. Lond. B Biol. Sci. 369, 20130440.

Brandvain Y, Barker MS, Wade MJ. 2007. Gene co-inheritance and gene transfer. Science 325(5819), 1685.

Brandvain Y, Wade MJ. 2009. The functional transfer of genes from the mitochondria to the nucleus: the effects of selection, mutation, population size and rate of self-fertilization. Genetics 182, 1129–1139.

Burton RS, Ellison CK, Harrison JS. 2006. The sorry state of F2 hybrids: consequences of rapid mitochondrial DNA evolution in populations. American Naturalist 168, S14–S24.

Clancy DJ, Hime GR, Shirras AD. 2011. Cytoplasmic male sterility in Drosophila melanogaster associated with a mitochondrial CYTB variant. Heredity 107, 374–376.

Dean R, Zimmer F, Mank JE. 2014. The potential role of sexual conflict and sexual selection in shaping the genomic distribution of mito-nuclear genes. Genome Evol. Biol. 6(5), 1096–1104.

Dobler R, Dowling DK, Morrow EH, Reinhardt K. 2018. A systematic review and meta-analysis reveals pervasive effects of germline mitochondrial replacement on components of health. Human Reproduction Update 24(5), 519–534.

Dowling DK, Abiega KC, Arnqvist G. 2007. Temperature-specific outcomes of cytoplasmic-nuclear interactions on egg-to-adult development time in seed beetles. Evolution 61, 194–201.

Dowling DK and Adrian RE. 2019. Challenges and prospects for testing the mothers curse hypothesis. Integrative and comparative biology 59 (4), 875–889.

Drown DM, Preuss KM, Wade MJ. 2012. Evidence of a Paucity of Genes That Interact with the Mitochondrion on the X in Mammals. Genome Biol. Evol. 4(8), 875–880.

Frank SA, Hurst LD. 1996. Mitochondria and male disease. Nature 383, 224.

Gallach M, Chandrasekaran C, Betran E. 2010. Analyses of nuclearly encoded mitochondrial genes suggest gene duplication as a mechanism for resolving intralocus sexually antagonistic conflict in Drosophila. Genome Biol. Evol. 2, 835–850.

Gemmell NJ, Metcalf VJ, Allendorf FW. 2004. Mother’s curse: the effect of mtDNA on individual fitness and population viability. Trends Ecol. Evol. 19, 238–244.

Giordano L, Sillo F, Gonthier P. 2018. Mitonuclear interactions may contribute to fitness of fungal hybrids. Sci Reports 8, 1706.

Greiner S, Sobanski J, Bock R. 2015. Why are most organelle genomes transmitted maternally? Bioessays 37, 80–94.

Hadjivasiliou Z, Lane N, Seymour RM, Pomiankowski A. 2013. Dynamics of mitochondrial inheritance in the evolution of binary mating types and two sexes. Proc R Soc B 280(1769), 20131920.

Havird JC, Hall MD, Dowling DK. 2015. The evolution of sex: A new hypothesis based on mitochondrial mutational erosion. BioEssays 37(9), 951–958.

Johnston IG, Burgstaller JP. 2019. Evolving mtDNA populations within cells. Biochem. Soc. Trans. 47 (5), 1367–1382.

Keaney TA, Wong HWS, Dowling DK, Jones TM, Holman L. 2019. Mother’s curse and indirect genetic effects: Do males matter to mitochondrial genome evolution? Journal of Evolutionary Biology 33(2), 189–201.

Kondrashov AS. 1993. Classification of Hypotheses on the Advantage of Amphimixis. Journal of Heredity 84(5), 372–387.

Kuijper B, Lane N, Pomiankowski A. 2015. Can paternal leakage maintain sexually antagonistic polymorphism in the cytoplasm? Journal of Evolutionary Biology 28(2), 468–480.

Lane N, Martin W. 2010. The energetics of genome complexity. Nature 467(7318), 929.

Lane N. 2011. Energetics and genetics across the prokaryote-eukaryote divide. Biol. Direct 6(1), 35.

Ma H, Gutierrez NM, Morey R, et al. 2016. Incompatibility between Nuclear and Mitochondrial Genomes Contributes to an Interspecies Reproductive Barrier. Cell Metabolism 24(2), 283–294.

Malik SB, Pightling AW, Stefaniak LM, Schurko AM, Logsdon Jr. JM. 2008. An Expanded Inventory of Conserved Meiotic Genes Provides Evidence for Sex in Trichomonas vaginalis. PLoS ONE 3(8), e2879.

Martin W. 1998. The hydrogen hypothesis for the first eukaryote. Nature 392(6671), 37–41.

Muller HJ. 1964. The relation of recombination to mutational advance. Mutation Research 1, 2–9.

McKenzie M, Lazarou M, Thorburn DR, Ryan MT. 2007. Analysis of mitochondrial subunit assembly into respiratory chain complexes using blue native polyacrylamide gel electrophoresis. Anal. Biochem. 364, 128–137.

Otto S. 2007. Unravelling the evolutionary advantage of sex: a commentary on “Mutation-selection balance and the evolutionary advantage of sex and recombination” by Brian Charlesworth. Genet. Res. 89, 447–449.

Radzvilavicius A. 2016. Evolutionary dynamics of cytoplasmic segregation and fusion: mitochondrial mixing facilitated the evolution of sex at the origin of eukaryotes. J Theor Biol 404, 160–168.

Radzvilavicius A, Lane N, Pomiankowski A. 2017. Sexual conflict explains the extraordinary diversity of mechanisms regulating uniparental inheritance. BMC Biology 15, 94.

Ramesh MA, Malik SB, Logsdon JM Jr. 2005. A phylogenomic inventory of meiotic genes; evidence for sex in Giardia and an early eukaryotic origin of meiosis. Curr Biol. 15(2), 185–191.

Rand DM, Haney RA, Fry AJ. 2004. Cytonuclear coevolution: the genomics of cooperation. Trends in Ecology & Evolution 19(12), 645–653.

Reinhardt K, Dowling DK, Morrow EH. Mitochondrial Replacement, Evolution, and the Clinic. Science 341(6152), 1345–1346.

Rogell B, Dean R, Lemos B, Dowling DK. 2014. Mito-nuclear interactions as drivers of gene movement on and off the X-chromosome. BMC Genomics 15, 330.

Rossignol R, Faustin B, Rocher C, Malgat M, Mazat JP, Letellier T. 2003. Mitochondrial threshold effects. Biochemical Journal 370(3), 751–762.

Vaught RC and Dowling DK. 2018. Maternal inheritance of mitochondria: implication for male fertility. Reproduction 155(4), R159–R168.

Wade MJ, Brandvain Y. Reversing mother’s curse: selection on male mitochondrial fitness effects. Evolution 63(4), 1084–1089.

Wolff JN, Nafisinia M, Sutovsky P, Ballard JWO. 2012. Paternal transmission of mitochondrial DNA as an integral part of mitochondrial inheritance in metapopulations of Drosophila simulans. Heredity 10, 57–62.

Wolff JN, Ladoukakis ED, Enriquez JA, Dowling DK. 2014. Mitonuclear interactions: evolutionary consequences over multiple biological scales. Phil. Trans. R. Soc. B 369, 20130443.

Woloszynska M. 2010. Heteroplasmy and stoichiometric complexity of plant mitochondrial genomes-though this be madness, yet there’s method in’t. Journal of Experimental Botany 61(3), 657–671.

Xu J, Li H. 2015. Current perspectives on mitochondrial inheritance in fungi. Cell Health and Cytoskeleton 7, 143.

